# Genome-resolved metagenomics reveals the potential for selection for antibiotic resistance due to metal pollution in soil microbial communities near a copper-nickel mine site in Botswana

**DOI:** 10.64898/2026.05.29.728080

**Authors:** Katlego P. P. Makale, Ioannis D. Kampouris, Venecio U. Ultra, Oakantse Dineo, Doreen Babin, Gaolathe Rantong

## Abstract

Waste mismanagement and metal pollution are high in countries in the global south and often create challenging, extreme conditions for soil microbial communities, which might lead to selection for antibiotic resistance. However, soils from polluted sites in Sub-Saharan areas are rarely sampled. Thus, it remains unclear whether the potential for dissemination of antibiotic resistance exists in these areas. In the present study, we performed genome-resolved metagenomics on samples across a gradient around an area impacted by a copper-nickel mine in Botswana to identify the potential for co-selection for antibiotic resistance due to mine pollution. Specifically, we sampled over a gradient across a mine area in Botswana and performed genome-resolved metagenomics. In general, pollution increased with proximity to the mine, while prokaryotic diversity increased with increasing distance from the mine, and we could identify a few taxa that significantly correlated with the metal concentrations. Moreover, our assembled contigs indicate a potential for co-selection due to the dual function of these genes as both antibiotic resistance genes (ARGs) and metal resistance genes (MRGs), as they act as multidrug efflux pumps. In contrast, some ARGs co-occurred in the same region as MRGs, indicating the potential for co-selection due to co-localization. However, we could not detect any co-localization of ARGs/MRGs with horizontal gene transfer (HGT) markers at the contig level. We binned the contigs to MAGs, and we found several MAGs of high quality and completeness, belonging to taxa such as *Actinobacteriota, Chloroflexota, Acidobacteriota,* and *Dormibacterota,* that possess ARGs, MRGs, and HGT-markers. Moreover, we found ARGs, MRGs, and HGT-markers in eight MAGs, after binning, indicating no direct association of HGT-markers with ARG and MRG occurrence. In summary, our metagenomic analysis indicates that pollution in mines can lead to co-selection of AMR, specifically ARGs with a broad spectrum of substrates (e.g., efflux pumps) that can act as MRGs, in an under-sampled location such as Sub-Saharan countries.

**IMPORTANCE:** This study indicates the potential role of metal pollution from mismanaged mining activities in facilitating the selection for antibiotic resistance in soil microbial communities in a Sub-Saharan area located in Botswana. In combination with the lower amount of health regulations and hygiene standards, the dissemination of antibiotic resistance poses risks to human health through the environment in many Sub-Saharan countries. By performing genome-resolved metagenomics and identifying antibiotic resistance genes (ARGs) as metal resistance genes (MRGs) across diverse assembled genomes or co-localization, this work highlights the potential for antimicrobial resistance dissemination due to mine activities. Thus, these findings underscore the urgent need to understand the potential side effects of mine pollution on the spread of antimicrobial-resistant bacteria (ARB) and associated ARGs in Sub-Saharan countries.

## 1. Introduction

Dissemination of antibiotic resistance is considered a major global human threat, with millions of deaths per year, a number projected to increase in the following decades (Piddock 2012). Without a doubt, the consumption of antibiotics promotes the dissemination of antimicrobial resistance (AMR) in human and animal populations, as well as in the environment (Gillings et al. 2015; Piña et al. 2020; Kampouris et al. 2022; Soufi et al. 2025). Soil microbiomes drive nutrient cycling in terrestrial environments (Bahram et al. 2018; Cowan et al. 2022), determine soil fertility (Chaparro et al. 2012; Delitte et al. 2021) and health (Dubey et al. 2019). In addition, soil microbiomes also act as a reservoir of antibiotic resistance genes (ARGs) (Bahram et al. 2018). Specifically, most antibiotics have been discovered by studying antagonistic activities from soil microbes (Nesme and Simonet 2015). Thus, soil environments can act as a source of AMR dissemination via ARGs for old and new antibiotics, which might be selected by anthropogenic pressure (Nesme and Simonet 2015; Bahram et al. 2018; Manaia et al. 2018), like those induced by heavy metal contamination from mining activities.

Wastewater discharge and manure fertilization have been considered as the major sources of AMR pollution (Walters et al. 2015; Caucci et al. 2016; Karkman et al. 2019; Kampouris et al. 2021; De La Cruz Barron et al. 2023). However, apart from the direct release of ARG-carrying bacteria due to wastewater discharge or manure amendment (Binh et al., 2008; Cerqueira et al., 2019a, 2019b; Heuer et al., 2011; Heuer and Smalla, 2007), other anthropogenic activities might lead to the dissemination of AMR as well (Seiler and Berendonk 2012). Specifically, AMR dissemination might also be enhanced due to pressure from non-antibiotic compounds (Wang et al. 2019) or toxic metals (Zhao et al. 2019), which may lead to co-selection or increased horizontal gene transfer (HGT) events between microbes (European Crop Protection Association 2010; Seiler and Berendonk 2012; Zhang et al. 2018). Global efforts have been made to investigate the taxonomic composition of microbial communities across the world and their link to AMR (Bahram et al. 2018; Gharechahi et al. 2021; Cowan et al. 2022). However, there is limited information about ARG prevalence in African soil environments, especially those affected by environmental pollution (Cowan et al. 2022).

Several metals and toxic compounds have been shown to drive AMR dissemination through co-selection of ARGs, such as copper, which can promote AMR dissemination in controlled settings (Zhang et al. 2019). Since mining activity releases several metals and metalloids (Nagajyoti et al. 2010; Ramkrishna et al. 2016), these metals and metalloids may co-select for ARGs across different environments (Seiler and Berendonk 2012). In addition, multidrug efflux pump genes might have a broad range of substrates, conferring resistance to both toxic metals and a variety of antibiotics (Adhikary et al. 2022). Thus, unsurprisingly, some multidrug resistance genes encoding efflux pumps might be present in both ARG (Alcock et al. 2023) and metal resistance gene (MRG) databases (Pal et al. 2014).

Mining activities pose a major health threat in Africa, particularly in mining areas of countries with low GDP, such as Sub-Saharan Africa (SSA) (Takam Tiamgne et al. 2022). The absence of high health and environmental standards exacerbates the impact compared to countries with high GDP (Yameogo et al. 2021; Manyiwa et al. 2022). Due to mismanagement, metal pollution is high in these areas (Manyiwa et al. 2022), thereby, several mining areas in the SSA region create a challenging and extreme habitat for microbial communities (Oyetibo et al. 2021). However, there is no evidence that metal pollution could contribute to AMR dissemination in soil microbial communities in SSA countries. Botswana is a middle-income country whose economic growth is dependent on mining (Motlogelwa and Leeuw, 2023; Patel *et al*, 2021). Mining contributes 40% to the GDP, and some of the minerals produced include diamond, gold, copper, and nickel (Phiri *et al*., 2021). One of the mines in Botswana, the Bamangwato Concession Limited (BCL) Copper-Nickel mine in Selibe Phikwe, located between longitudes 27LJ47E and 27LJ53E and latitudes 22LJ55S and 22LJ00S, was operational for 42 years between 1974 and 2016. During its activity, the mine released pollutants such as heavy metals (including lead, arsenic, manganese, copper, and nickel), acid mine water, and air particulate matter, which have spread into the environment, affecting plants, microorganisms, animals, and potentially humans (Tchounwou *et al*., 2012; Gabasiane *et al*., 2019). Such a setting can provide further insights into the co-localizations of ARGs and MRGs in soil microbiota.

In the present study, we investigated soil microbial communities and their functional potential in a mining area in Botswana, specifically the BCL mine in Selibe Phikwe, using genome-resolved metagenomics. Thus, we aimed to evaluate whether metal pollution from an inactive mine site could contribute to AMR dissemination via co-localization of ARGs with MRGs and HGT-associated markers, using genome-resolved metagenomics. Specifically, we evaluated whether the pollution due to former BCL mining activity can potentially select for ARGs by performing deep shotgun sequencing of soils sampled across a gradient around the BCL mine. Overall, our study provides further insights into the occurrence of AMR and metal resistance in soil microorganisms from such a unique and under-investigated location.

## 2. Materials and Methods

### 2.1 Soil Sampling

The Selebi-Phikwe area features a Savanna-type vegetation (Fig. 1A) dominated by tree species such as *Colophospermum mopane* and *Vachellia* species, shrubs such as *Grewia* species, and grasses such as *Aristida congesta* (Vurayai et al. 2015; Manyiwa and Ultra 2022). Other grasses present include *Pennisetum setaceum, Stipagrostis uniplumis,* and *Cynodon dactylon,* and trees include *Boscia species, Senegalia nigrescens, Dichrostachys cinerea,* and *Faidherbia albida.* The vegetation is also made of several herbs, forbs, and sedges (Manyiwa et al. 2021).

**Figure 1:**
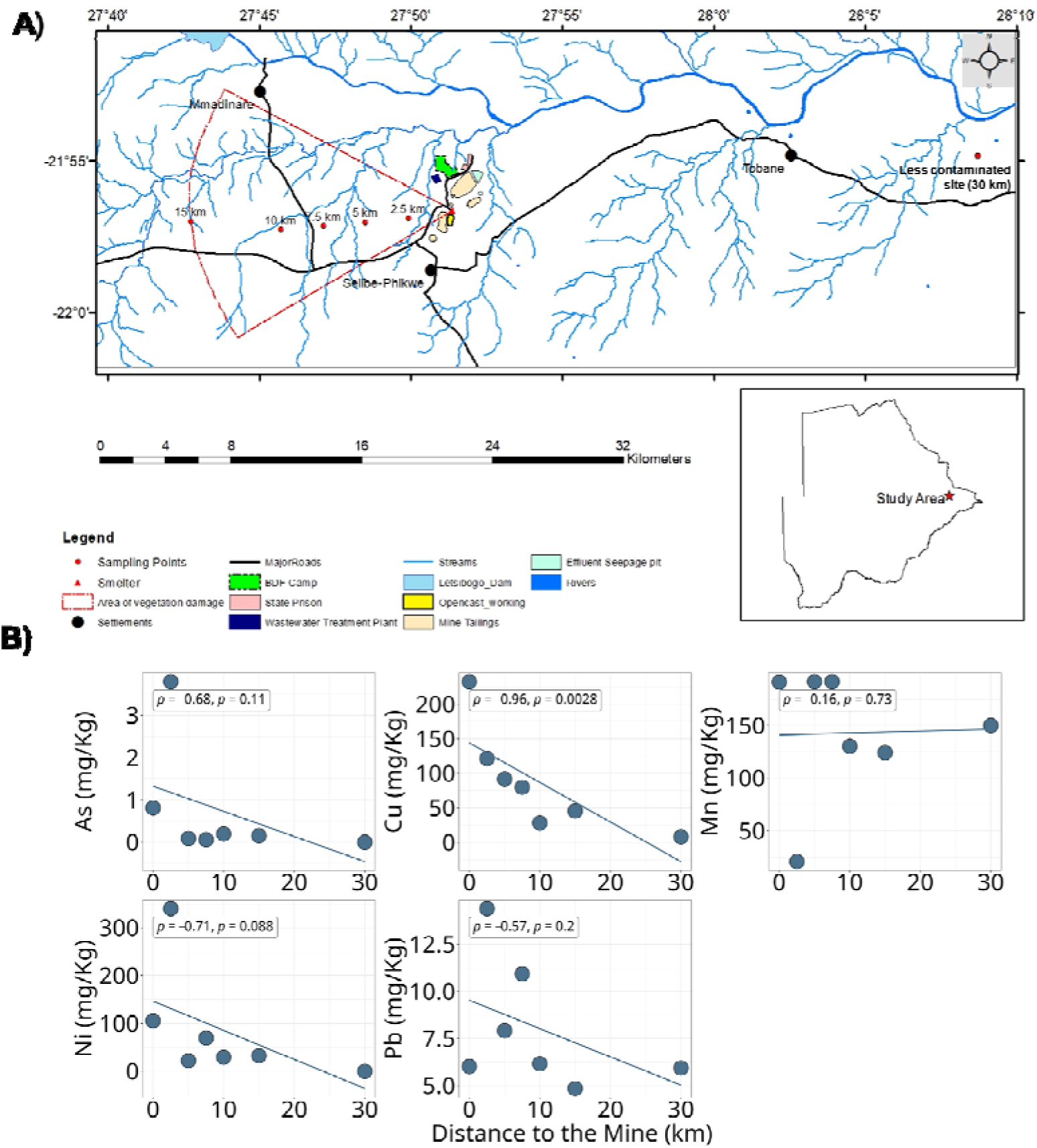
A) Map and location of sampled area (Selibe-Phikwe) along with description of mining activity. B) Regressions with the distance to the mine (smelter) and the concentration of various metals (Spearman rank correlation, n = 7).

Sites near the mine exhibit clear signs of damage, such as a lack of vegetation and sediment accumulation in water channels. Damage from mining activity appears to decrease with increasing distance and may also be influenced by wind drift direction. The soil samples were collected from seven different sites around the BCL mine. Six of the samples (0 km, 2.5 km, 5 km, 7.5 km, 10 km, and 15 km from the smelter, Fig. 1A) were collected on the western side of the smelter. The west side of the smelter is the windward (direction where the wind blows) and is the most contaminated side of the mine (Fig. 1A). In addition, one sample from a distant and less-polluted area was collected (30 km on the eastern side of the smelter, Fig. 1A). To collect soil samples, the zig-zag soil sampling method was used. Plant residue from the surface was removed, and a soil auger was used to collect the soil at 0-30 cm depth. Five soil samples were collected from each site to create a composite sample. The five samples were placed into a clean bucket, where stones were removed, and any large lumps were crushed to ensure the soil was loose and consistent. The soil was then thoroughly mixed and evenly spread in a container. After mixing, a composite sample was taken and transferred to a ziplock bag for transportation to the lab at a temperature of 4°C. Soil metal concentrations were analysed using a portable X-ray fluoresence (pXRF) (Delta-X, Massachusetts, USA).

### 2.2 DNA extraction

DNA was extracted directly from 250 mg of each soil sample using the ZYMO RESEARCH quick-DNA soil microbe mini-prep kit (Zymo Research, USA) and following the manufacturer’s protocol. The concentration of isolated DNA was measured using a NanoDrop (Thermo Fisher Scientific, USA) at 260 and 280 nm wavelengths. The genomic DNA was then run on a 1% agarose gel and visualized using an ultraviolet transilluminator.

### 2.3 Sequencing

The metagenomic sequencing was performed at the Agricultural Research Council (ARC) in South Africa. Briefly, the DNA libraries were prepared using the MGIEasy Universal DNA Library Prep Set (MGI Tech Co., Ltd., Shenzhen, China; Cat. Nos. 1000006985/6986/17571) following the manufacturer’s instructions. DNA was fragmented using a Covaris Focused-ultrasonicator (Thermo Fisher Scientific, USA), quantified with a Qubit™ 3 Fluorometer and dsDNA HS Assay Kit (Invitrogen, USA), and fragment size distribution was assessed using an Agilent 2100 Bioanalyzer (Agilent Technologies, USA). Library preparation included end repair, A-tailing, adapter ligation, bead cleanup, PCR amplification, and single-strand circularization. Sequencing was performed on the Illumina HiSeq 3000 platform (Illumina, San Diego, USA) using paired-end (PE100/PE150) chemistry.

### 2.4 Bioinformatics and sequencing analysis

Paired-end sequences were merged using PEAR (v0.9.11) (Zhang et al. 2014). PhixA genomes were removed using BBDuk from BBTools (v39.12) (Bushnell et al. 2017). Prokaryote abundance was evaluated with SingleM (v0.18.3) (Woodcroft et al. 2024), which takes into account single-copy marker genes and their coverage. Sequences were assembled using the MEGAHIT algorithm (v1.2.9) (Li et al. 2015) with a stepwise approach starting from 17 kmers and reaching 127 kmers (--k-min 17-- k-max 127-- k-step 10). Minimum contig length was set at 1500 bp. Following the assembly, the contigs were mapped to the reads using NGLess (v.1.5) (Coelho et al. 2019) and BWA-MEM (v.0.7.17) (Li and Durbin 2010). The output of the contigs and the mapped read coverage were used to generate metagenomics bins using the algorithm SemBin2 (Pan et al. 2023). Bin completeness and contamination were inferred by the CheckM2 algorithm (Chklovski et al. 2023) based on the lineage of single-copy genes. Bins were classified using the genome taxonomy database (GTDB) taxonomy classifier (GTDB-tk) (Parks et al. 2022) (Chaumeil et al. 2020). For antibiotic resistance gene (ARG) annotation, we mapped bins to the Comprehensive Antibiotic Resistance Database (CARD, v4.0.0), which includes intrinsic and horizontally transferred ARGs (Alcock et al. 2023). To annotate metal resistance genes (MRGs), we used the predicted MRG database from BacMet (v2.0.0) (Pal et al. 2014). For annotating horizontal gene transfer (HGT) markers, we used the HGT-marker database (Pärnänen et al. 2018). Every database was translated into amino acids from nucleotides using EMBOSS transeq (v6.60) (McWilliam et al. 2013). Initial gene annotation and prediction of proteins were performed with Prokka (Seemann 2014). Translated genes to proteins were annotated for each type of database using DIAMOND aligner (Buchfink et al. 2014), with 70% identity, minimum alignment of 30 amino acids, and the default e-value cutoff of DIAMOND (blastp command). All data were analyzed in R (v4.3.0) (R Core Team 2021) using the packages tidyverse (v2.00) (Wickham et al. 2019), ggplot2 (3.4.3) (Hadley Wickham 2016), dplyr (Wickham et al. 2023), ggpubr (Kassambara 2023), stringr (Wickham 2023), and reshape2 (v1.4.4) (Wickham 2007). Richness and Shannon-Index (α-diversity) were evaluated using the “vegan” R package (Oksanen et al. 2025). Correlations were performed with Spearman’s rank correlation, and *p*-values were adjusted for multiple testing with Benjamini-Hochberg correction. Scripts were uploaded to the https://github.com/JonKampouris/BCL_Mine_Soil_Metagenomics repository, and sequencing data were uploaded to SRA with BioProject accession PRJNA1332977. Scripts for the data analysis can be found in the following repository https://github.com/JonKampouris/BCL_Mine_Metagenomes.

## Results

### Distance to the smelter negatively correlates with copper concentrations

To estimate whether the distance to the smelter correlates with increased pollution due to mismanagement, we evaluated whether the increased metal concentrations correlated with the distance to the smelter by performing Spearman’s rank correlation analyses (Fig. 1B). Out of all metals, copper showed a significant negative correlation with increasing distance from the smelter (Spearman’s rank ρ = −0.96, *p* = 0.0028, n = 7, Table 1). Other metals, such as nickel and arsenic, also negatively correlated with the increasing distance to the smelter, but this correlation was not significant (Spearman’s rank ρ < −0.5, *p* > 0.05, n = 7, Table 1). This indicated that the mining activity uniformly distributed copper concentrations as a gradient across the area since copper concentration significantly increased as a function of proximity to the smelter area.

**Table 1.** The texture of soils from sampled sites in the area near the smelter from the BCL mine.

| **Sites and distance**<br>**from the mines** | **As (mg Kg<sup>-1</sup>)** | **Cu (mg Kg<sup>-1</sup>)** | **Mn (mg Kg<sup>-1</sup>)** | **Ni (mg/Kg)** | **Pb (mg/Kg)** |
| --- | --- | --- | --- | --- | --- |
| Site1 (0 km) | 0.8114 | 232.8 | 191.32 | 105.74 | 6.022 |
| Site2 (2.5 km) | 3.796 | 121.5 | 20.76 | 339.8 | 14.392 |
| Site 3 (5 km) | 0.08874 | 91.66 | 191.7 | 22.36 | 7.924 |
| Site 4 (7.5 km) | 0.061 | 79.74 | 191.7 | 70.02 | 10.938 |
| Site 5 (10 km) | 0.19778 | 28.2 | 130.3 | 29.7 | 6.164 |
| Site 6 (15 km) | 0.15426 | 45.26 | 124.3 | 32.82 | 4.848 |
| Site 7 (30 km) | 0 | 8 | 150 | 0 | 5.94 |

### Distance to the smelter positively correlates with prokaryote diversity and strongly shapes prokaryote community composition

To verify the effects of the distance to the smelter on prokaryotic diversity, we assessed prokaryotic diversity using an approach based on several single-copy marker genes (SingleM). In general, the distance from the smelter positively correlated with the Shannon-Index, indicating that metal pollution impacted soil α-diversity, leading to the prevalence of specific taxa (Fig. 2A, ρ = 0.86, *p* = 0.024, n = 7). In addition, we evaluated which taxa correlate with distance from the smelter: overall, we found only one taxon classified as *JADMIN01* family (Phylum: *Chloroflexota*, Class: *Ktedonobaceria*) to negatively correlate with the distance to the smelter (Spearman’s rank ρ = −0.99, adj. *p*-value = 0.002, n=7), indicating that the metal pollution selects for this taxon. In contrast, other *Chloroflexota* members (Class: *UBA677,* Fig. 2B) were positively correlated to the distance to the smelter (Spearman’s rank ρ > 0.9, adj. *p*-value < 0.05, n=7). However, due to the low number of samples, we found only a few correlations that indicate that metal pollution decreased or increased the relative abundance of several taxa. In conclusion, proximity to the smelter is connected with higher metal concentrations, shapes community composition, and decreases α-diversity.

**Figure 2.**
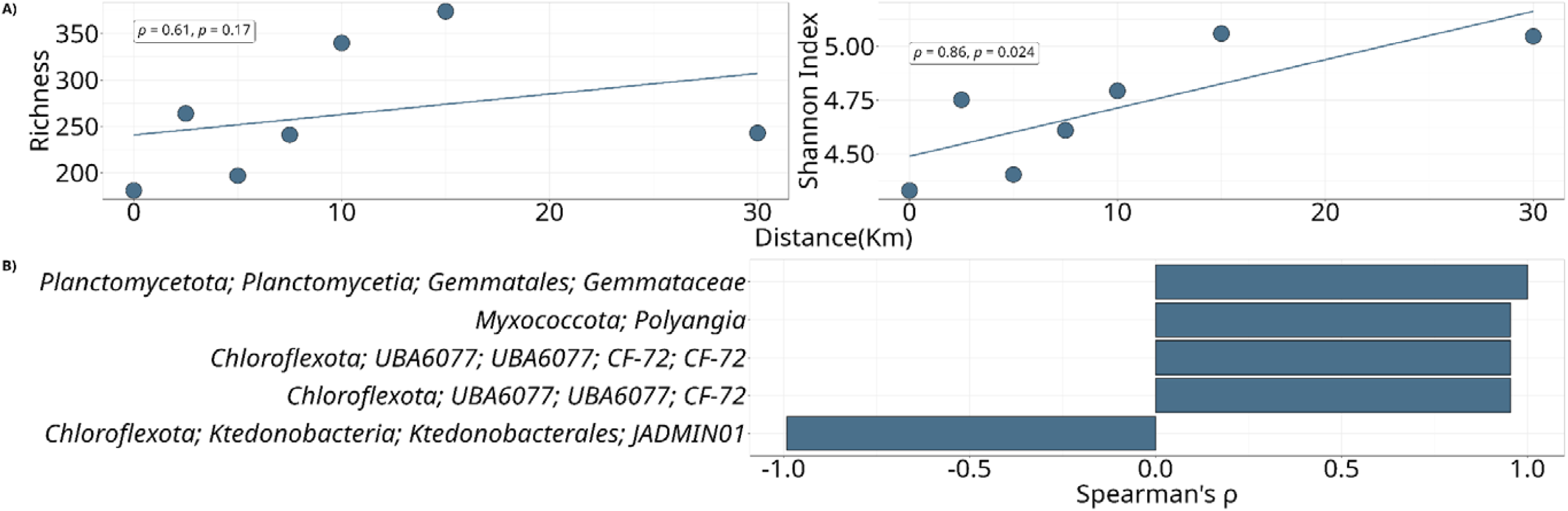
A) Regression of distance to the smelter (km) and α-diversity, based on read abundance of OTUs inferred by the singleM algorithm that utilizes single-copy marker genes for calculating the relative abundance of taxa. B) SingleM taxa that correlate with the distance to the smelter. SingleM OTUs were grouped until genus level (or at the lowest taxonomy level), and Spearman’s correlation with *p*-value correction was performed with Benjamini-Hochberg correction. Only taxa with adjusted *p*-value < 0.05 are shown.

### Potential for co-selection of ARGs with MRGs based on contig-level read assemblies

To investigate the potential occurrence of ARGs, MRG, and HGT-markers in prokaryotic genomes from the mine area, we performed assembly of the metagenomic reads. Our assembly generated 174686 contigs of 602188885 bp total length, with 24955.14286 ± 11036.73744 contigs per sample (mean ± standard deviation). Generally, we found annotated MRGs and ARGs in the herein analyzed contigs, and examined whether these genes were co-localized or annotated as both ARGs and MRGs (Fig. 3, Table S1, S2, and S3). Interestingly, several contigs contained MRGs that conferred resistance to copper, such as *copF* and *copR* (Table S1). In most cases (n=16), these annotated ARGs or MRGs were the same genes (or similar gene variants) that were present in both databases, and thus were annotated for both resistance to metals, but also resistance to antibiotics (e.g., *acrB*, *acrD*, and *acrF* variants), at the same contig locus (Fig. 3, Table S1 & S2). These genes encode efflux pumps that confer resistance to antibiotics, disinfectants, and metals (Table S1 & S2). Meanwhile, we found both MRG and ARG co-localization in a different locus of the same contig, such as co-localization of the MRG *cpx*R and the ARG *cpx*A, for eight contigs (Fig. 3, Table S1 & S2). However, we did not find any HGT-marker gene co-localized in the contigs with either ARGs or MRGs (Table S1 & S2). In conclusion, the contig-level assemblies indicate the potential for co-selection for AMR due to metal pollution, mostly due to genes with high substrate affinity, such as multidrug efflux pump, which can target both ARGs and MRGs (Fig. 3, Table S1 & S2), while co-localization of different genes in the same contig can occur as well.

**Figure 3.**
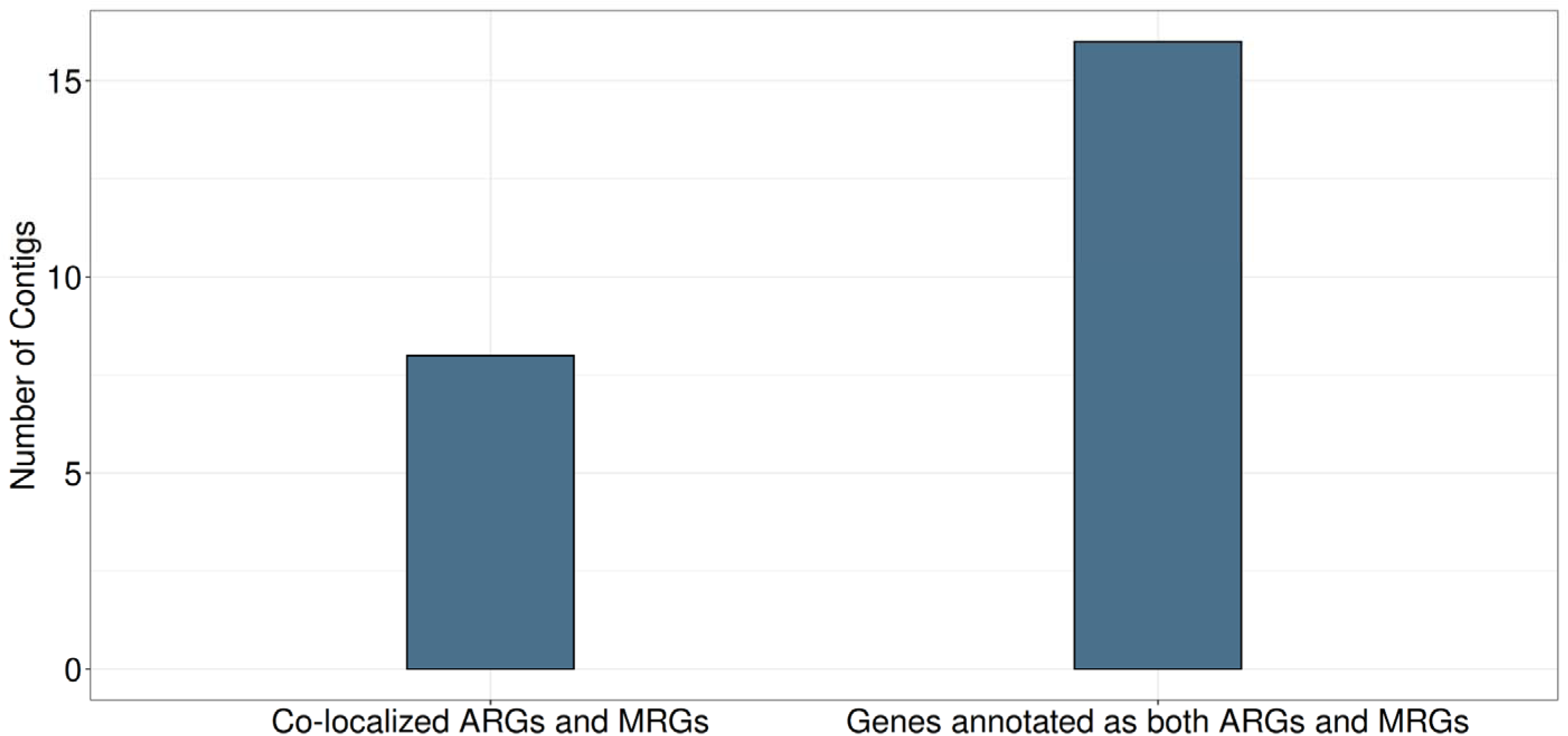
Number of assembled contigs with annotated metal resistance gene (MRGs) and antibiotic resistance genes (ARGs) using the BacMetPred and CARD databases, respectively. Contigs were screened for co-localization (different mapped regions and annotation due to bifunctionality of one gene as both ARG and MRG (e.g., efflux pumps).

### Generation of metagenome-assembled genomes (MAGs) and their taxonomy

To investigate the occurrence of ARGs, MRG, and HGT-markers in prokaryotic genomes from the mine area, we performed binning of the assembled metagenomic contigs. Our assembly generated 253 metagenome assembled genomes (MAGs) using MEGAHIT and SemiBin2, respectively (Fig. 4, Table S4). In total, 55 MAGs showed quality above 50% and contamination with less than 5%. Moreover, 35 MAGs showed a higher quality of above 80% with less than 5% contamination (High Quality MAGs, Fig. 4, Table S4). These high-quality MAGs were classified as *Acidobacteriota (Acidobacteria*)*, Thermoproteota* (*Nitrosphareota*)*, Eremiobacterota, Dormibacterota, Chloroflexota* (*Chloroflexi*), and *Actinomycetota* (*Actinobacteria*) (Fig. 4, Table S4).

**Figure 4.**
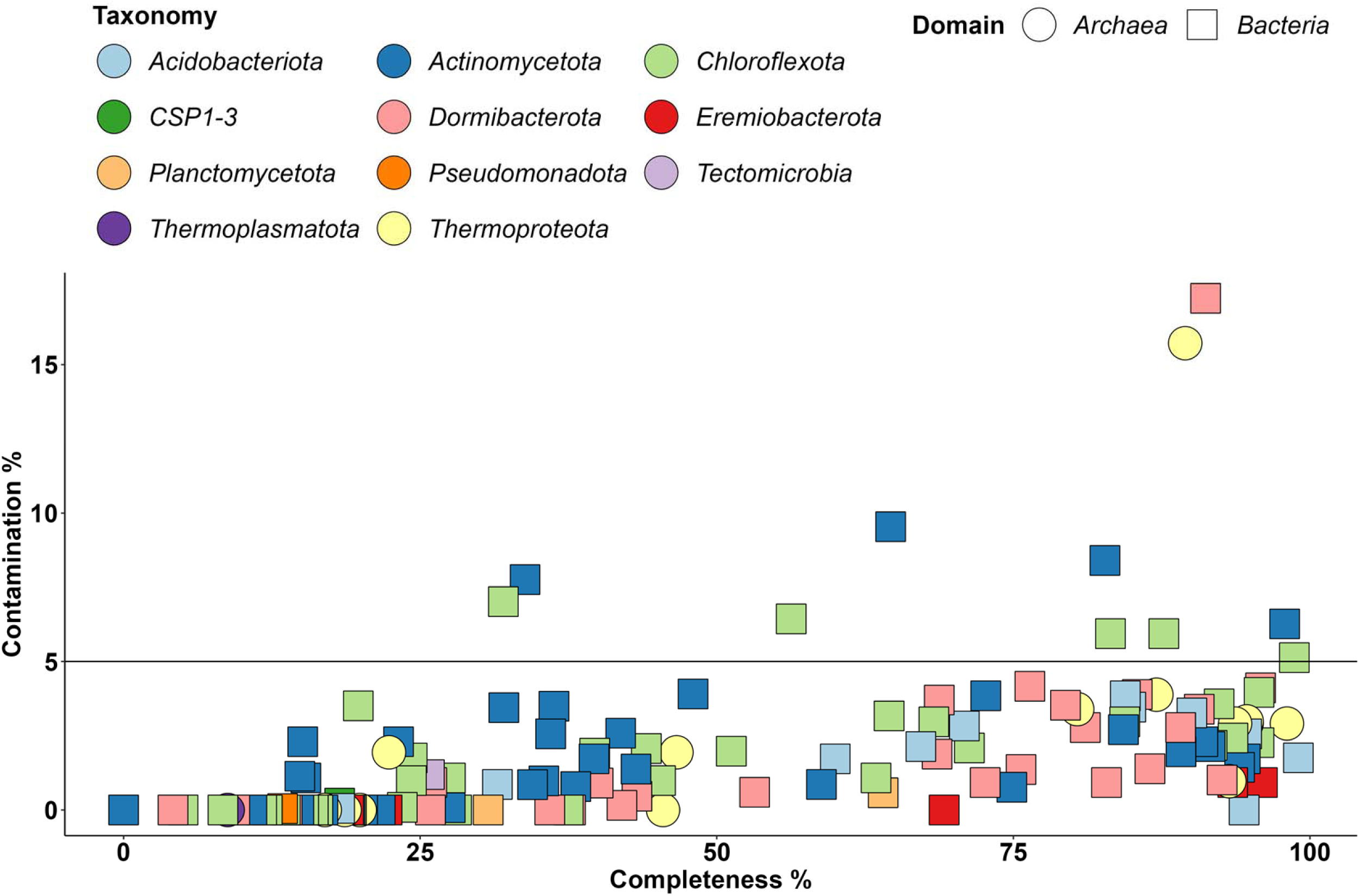
Completeness and Contamination of Metagenome assembled genomes and their taxonomy at Phylum level.

### Presence of antibiotic resistance genes in metagenome-assembled genomes

We found ARG presence in 132 out of the 253 MAGs. MAGs carrying resistance determinants were classified as *Acidobacteriota*, *Chloroflexota*, *Actinomycetota*, *Eremiobacterota*, *Dormibacterota*, *Planctomycetota*, *Tectomicrobia*, CSP1-3, and *Pseudomonadota* (Fig. 5). In general, *Actinomycetota* contain ARGs resistant to various classes of antibiotics such as aminoglycosides and erythromycin. *Chloroflexota* and *Dormibacterota* MAGs also contained resistance genes to several classes of antibiotics (Fig. 5, Table S5). Surprisingly, several ARGs were annotated in MAGs with low completeness (13% complete), such as the *Pseudomonadota* MAG (Fig. 5, Table S5), which was classified as *Pantoea* and contained several genes that confer resistance to several classes of antibiotics. Overall, most annotated genes conferred resistance to elfamycin (n = 128), aminoglycoside (n = 82), glycopeptide (n = 24), and pyrazine (n = 24) antibiotics (Fig. 5, Table S5). In summary, we discovered the presence of several ARGs in MAGs belonging to different phyla.

**Figure 5.**
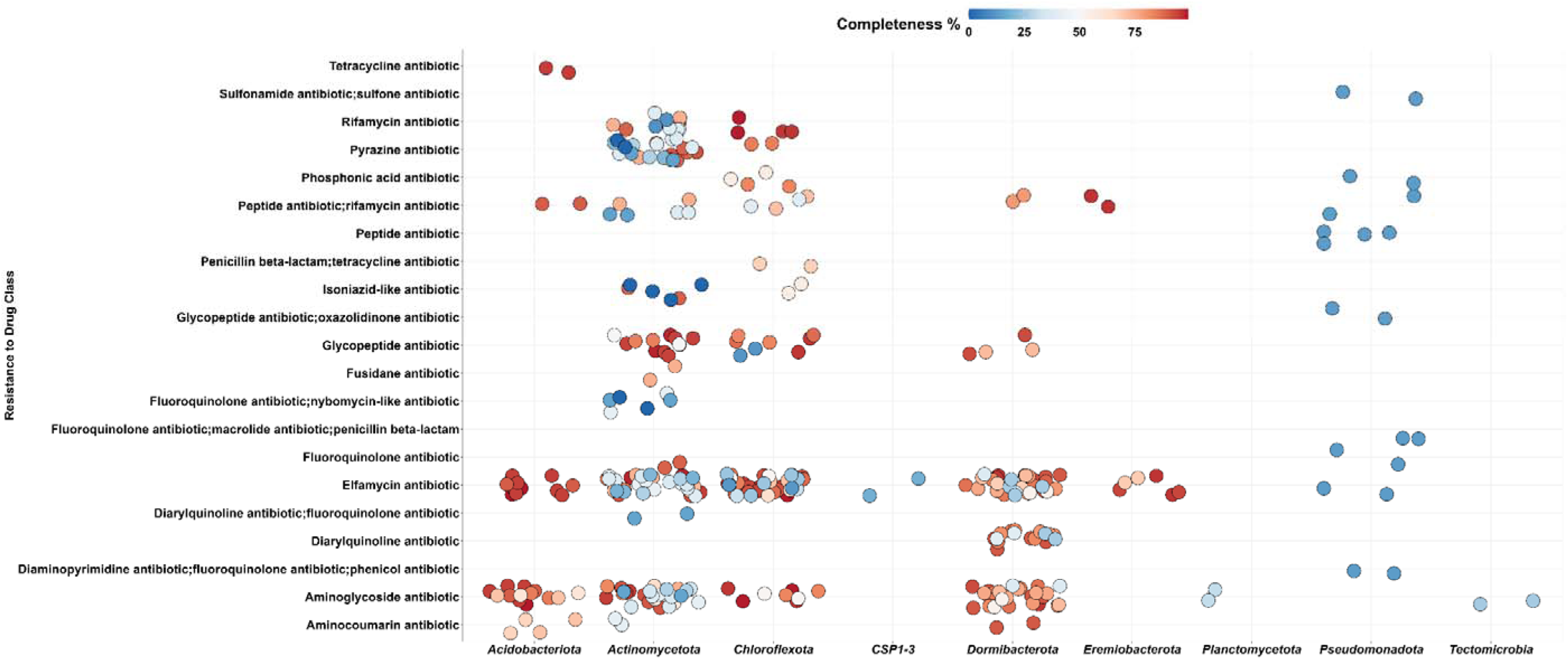
Class of annotated antibiotic resistance genes from the CARD database from several MAGs. The y-axis shows the drug class to which these genes confer resistance. The x-axis shows the classification at the Phylum level of the MAGs from which these genes were annotated. The color shows the completeness of the MAGs.

### 3.6 Presence of metal resistance genes in metagenome-assembled genomes

We found MRG presence in 63 out of the 253 MAGs. MAGs carrying MRG determinants were classified as *Acidobacteriota*, *Chloroflexota*, *Actinomycetota*, *Eremiobacterota*, *Dormibacterota*, *Pseudomonadota*, and *Thermoproteota* (*Archaea*) (Fig. 6, Table S6). We specifically focused on MRGs conferring resistance to As, Cu, and Ni, since these metals were found in high concentrations in the soil near the smelter area. As with ARGs, we found that MRGs were present in MAGs with low completeness, such as *Pseudomonadota* MAGs (*Pantoea*, Fig. 6). In general, most MAGs contained genes that confer resistance to As. However, *Actinomycetota* and *Chloroflexota* MAGs possessed putative MRGs to Cu (Fig. 6, Table S6). This is probably associated with selection for *Chloroflexota,* as a function of the distance to the smelter (Fig. 2B). Yet, other *Chloroflexota* taxa show an opposite relationship of the distance to the smelter (Fig. 2B). *Dormibacterota* genes were also annotated as MRGs to Cu and As (Fig. 6, Table S6). At the gene level, most of the genes (n=38) were annotated as *arsM*, which encodes arsenite methyl-transferase, eight genes were annotated as the copper resistance gene *copF,* and eight genes were annotated as *pstB* (Table S6). The rest of the MRG annotations occurred less than seven times. In summary, *Actinomycetota* and *Chloroflexota* MAGs contained more MRGs than the rest of the MAGs, and resistance to As, rather than Cu, was the most prevalent type of MRG class (Fig. 6, Table S6).

**Figure 6.**
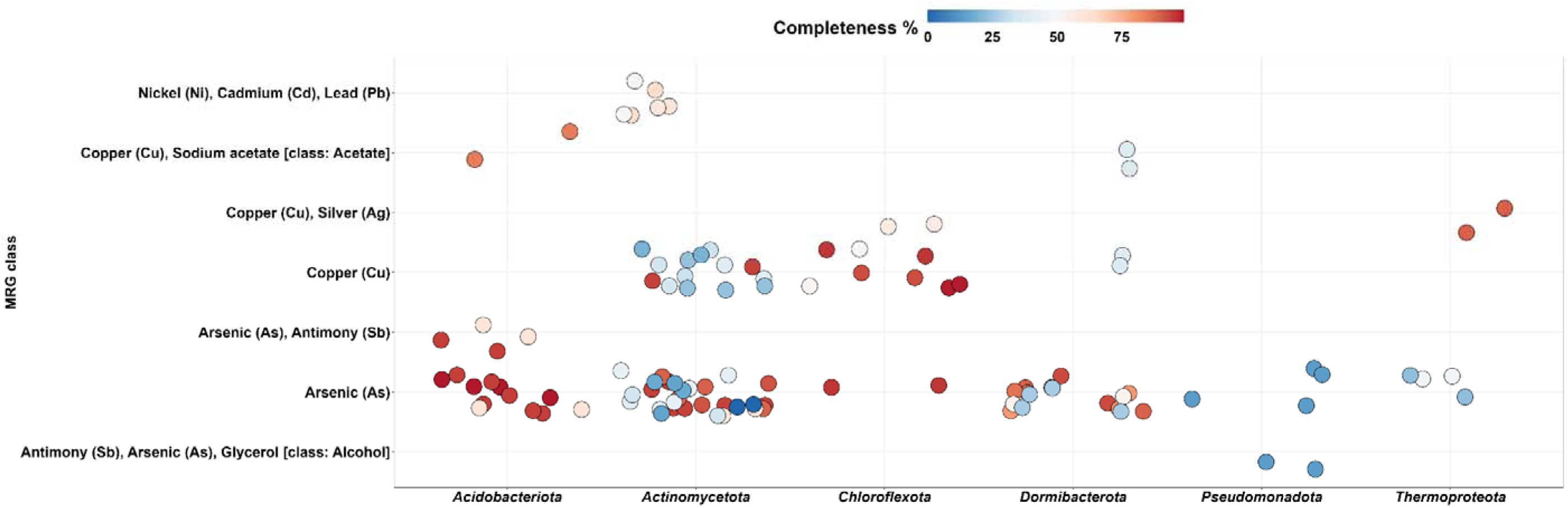
Class of annotated metal resistance genes (MRGs) from BacMetPred database. These genes were annotated from several MAGs, and one MAG might contain more than one gene. The y-axis shows the metal compound(s) that these genes confer resistance to specific metals and compounds (e.g., acetate) as defined in BacMetPred database. The x-axis shows the classification at the Phylum level of the MAGs from which these genes were annotated. The color shows the completeness of the MAGs from which these genes were annotated.

### 3.7 Horizontal gene transfer markers in metagenome-assembled genomes

Only 16 out of 253 MAGs contained HGT markers. These MAGs were classified as *Acidobacteriota, Actinomycetota, Chloroflexota, Dormibacterota,* and *Eremiobacterota* (Fig. 7, Table S7). Surprisingly, we found MAGs containing the gene *intI1* classified as *Acidobacteriota* and *Chloroflexota*. The transposase gene *tnpA* was detected across *Acidobacteriota*, *Actinomycetota*, *Chloroflexota*, and *Eremiobacterota* MAGs. The rest of HGT markers did not show such high prevalence across different phyla (Fig. 7, Table S7).

**Figure 7.**
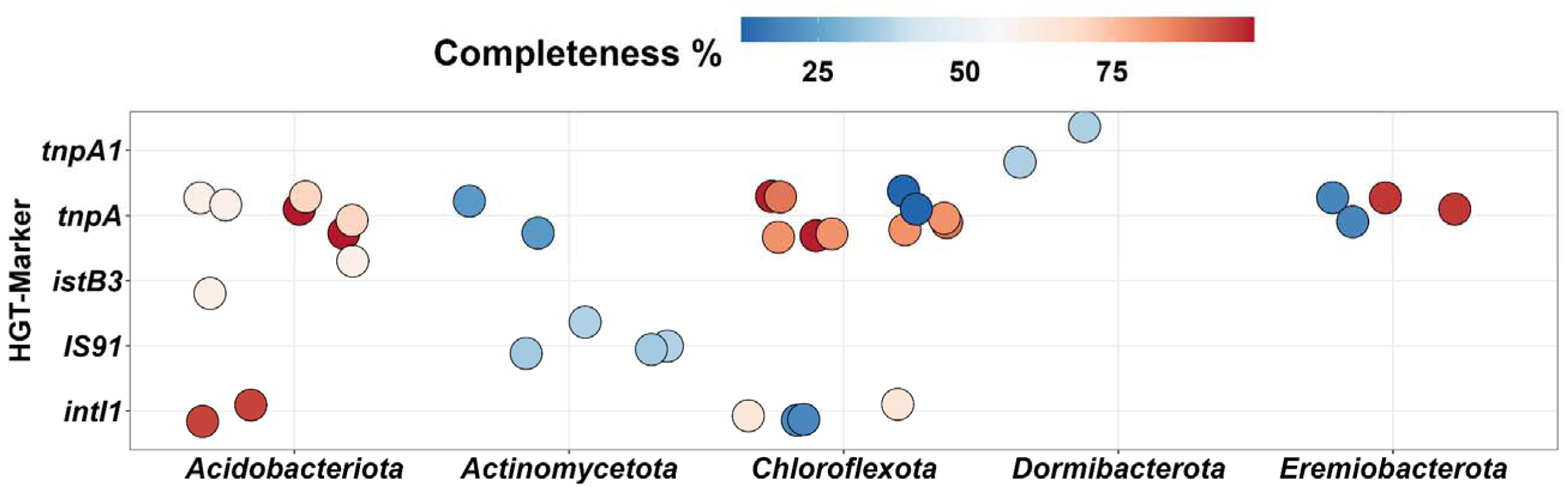
Annotated horizontal gene transfer (HGT) markers. These markers were annotated from several MAGs. The y-axis shows the metal compound(s) that these genes confer resistance to. The x-axis shows the classification at the Phylum level of the MAGs from which these genes were annotated. The color shows the completeness of the MAGs that were these genes annotated from.

### 3.8 Presence of antibiotic/metal resistance genes and HGT markers in the same MAGs

Binning revealed additional ARG, MRG, and HGT marker co-localization at the bin level (n = 6 MAGs, Fig. 8A). However, all HGT markers were co-localized in different contigs, since there was no co-localization of HGT markers with ARGs and MRGs at the contig level (Fig. 3). The MAGs having both HGT-markers, MRGs, and ARGs annotated were classified as *Chloroflexota, Acidobacteriota,* and *Actinomycetota*. In total, 38 MAGs were annotated for both ARGs and MRGs; eight MAGs showed both MRGs and HGT-markers, and eleven MAGs showed the presence of both ARGs, MRGs, and HGT-markers. Out of the six MAGs that were annotated for ARGs, MRGs, and HGT-markers, the annotated ARGs and MRGs were not variants of the same annotated gene located in both databases (Table S8). Thus, they were co-localized genes in the same bin. In addition, *intI1* was present in one *Acidobacteriota* MAG (Fig. 8B) that contained both ARGs and MRGs. In conclusion, binning grouped together HGT-markers and MRGs/ARGs, which were located in different contigs, but were part of the same bin (Fig. 8, Table S8).

**Figure 8.**
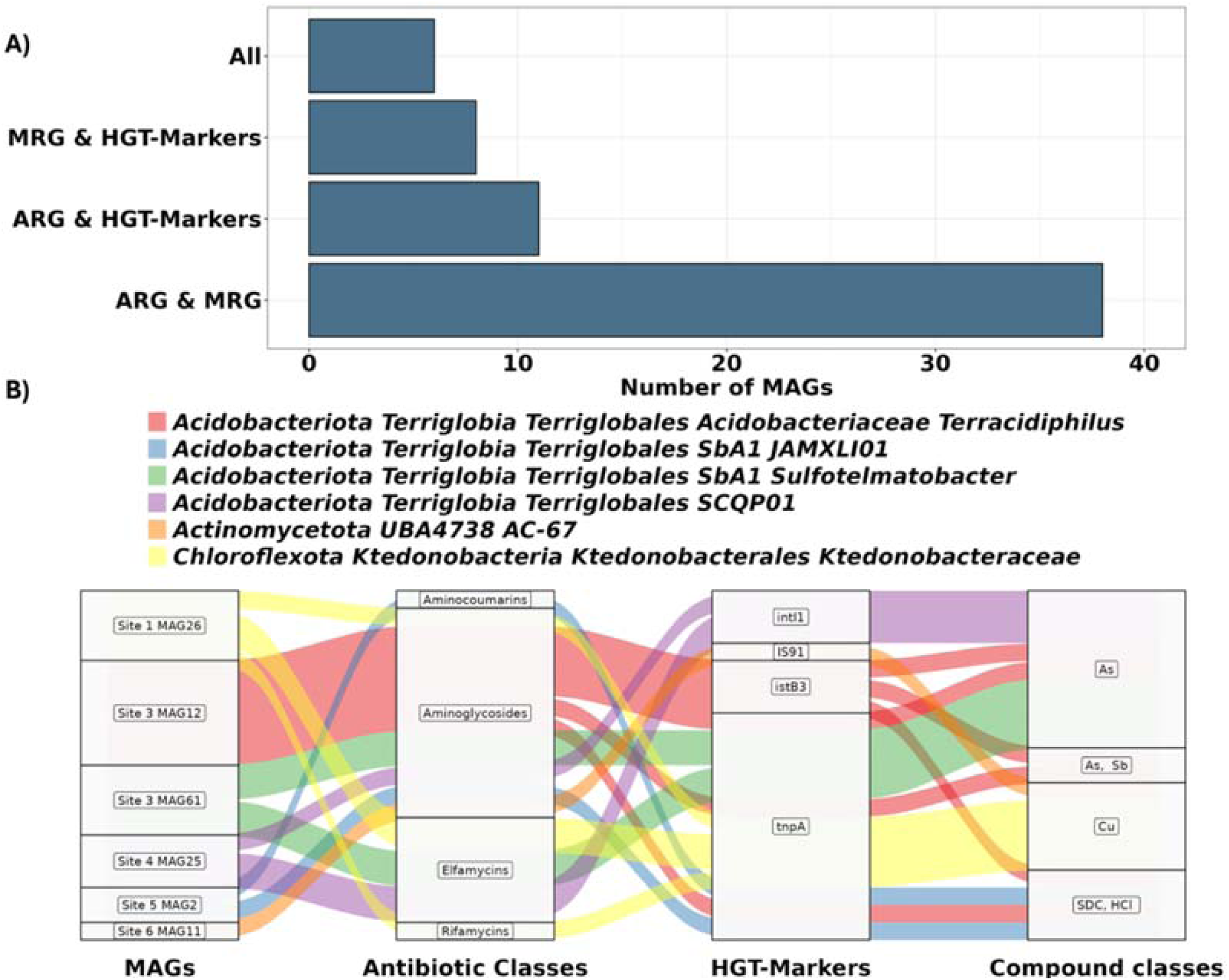
A) Number of metagenome-assembled genomes (MAGs) with annotated genes for antibiotic resistance genes (ARGs) and metal resistance genes (MRGs), along with horizontal gene transfer (HGT) markers. B) Alluvial plot of MAGs that contain the metagenome assembled genomes (MAGs), the antibiotic classes of the annotated ARGs, the annotated horizontal gene transfer markers (HGT markers), and the compound classes for the annotated metal resistance genes (MRGs). The color depicts the taxonomy of each MAG at the lowest possible level.

## 4. Discussion

Anthropogenic pollution, such as metal pollution, is known to cause toxicity to environmental microbes and has often been associated with selection for antimicrobial resistance (Seiler and Berendonk 2012; Zhang et al. 2018; Wang et al. 2021). While this selection presence has been extensively demonstrated in the lab (Zhang and Straight 2019; Zhang et al. 2019) we lack empirical evidence on whether this selection can be facilitated in areas located in developing countries, which are highly impacted by mining activities (Manyiwa et al. 2022; Gaothobogwe et al. 2025). Here, we analyzed seven soil samples using deep sequencing and metagenomic assembled genomes (MAGs). Our results provide empirical evidence that copper pollution reduced microbial diversity and increased one *Chloroflexota* taxon in an inactive mine located in Botswana. Since several ARGs were also annotated as MRGs due to widespread substrate targets, our genome-resolved metagenomic analysis indicated potential co-selection for metal resistance and AMR through multidrug efflux pumps. Moreover, only a few MAGs contained ARGs, MRGs, and HGT markers, but not in the same contigs, which indicated a low potential for HGT of these genes to other microbial taxa, under these conditions. In conclusion, using deep sequencing and genome-resolved metagenomics, we here provide evidence that metal pollution in an inactive, mismanaged mining site in Botswana can potentially promote the selection for AMR through the selection for ARGs with a broad spectrum of substrates, such as multidrug efflux pumps.

### 4.1. Selection for antibiotic resistance as a function of mine pollution

Metal pollution is considered to increase AMR dissemination through several mechanisms, such as selecting for multidrug resistance efflux pumps (Pal et al. 2014; Adhikary et al. 2022) and for microbial taxa that harbor both MRGs and ARGs (Seiler and Berendonk 2012). In addition, pollution is considered to cause stress to microbes, leading to energy-expensive HGT events, in order for the microorganisms to cope with the stress (Zhang et al. 2019). Using metagenome assembly and genome-resolved metagenomics, we here managed to confirm potential co-selection for AMR and metal resistance in Botswana, a country that is impacted by mine pollution due to mismanagement of mine activity (Manyiwa et al. 2022; Gaothobogwe et al. 2025). Proximity to the smelter increased metal pollution, and this pollution was associated with reduced microbial α-diversity in soil, even though the site was inactive. Thus, diversity dropped in occurrence with high stress for the microbial communities, in agreement with previous works (Rocca et al. 2019). This also coincides with the effect of the pollution on vegetation in the area near the mine (Letshwenyo 2016), indicating that the loss of diversity in the microbial fraction also coincides with the damage to overall diversity for local fauna. Often, environmental factors and pollution stress heavily shape the diversity and the composition of microorganisms (Yan et al. 2020). Despite the low number of samples, our correlation analysis indicated that metal pollution strongly shaped diversity and abundance. It is clear that these effects are indicative of the impact of such practices on the local soil microbiota. The examined mining has been inactive for several years, since 2016, indicating a surprising persistence of the effects that reduced diversity. Since plant rhizodeposition provides organic matter that maintains the function and abundance of soil microorganisms, we speculate that the loss of vegetation diversity might also contribute to the loss of microbial diversity. Thus, the impact of the mining on vegetation and soil microbiota might remain for several years.

Our deeply sequenced metagenomes managed to provide several assembled contigs and MAGs of different quality that contained genes that encoded efflux pumps that confer resistance to both metal and antibiotics, and were annotated using both CARD (antibiotics) (Alcock et al. 2023) and BacMet databases (metal resistance) (Pal et al. 2014). In addition, in a few cases, there was a co-localization of MRGs and ARGs in different loci of the same contig, such as the co-localization of the MRG *cpx*R and the ARG *cpx*A. Since *cpx*R is known to activate the expression of *cpx*A in bacterial cells facing stress (López et al. 2018) it is not surprising that these genes were found to be annotated in the same contig.

Furthermore, six MAGs contained ARGs, MRGs, and HGT-markers in different contigs of the same bins (Fig. 8). The presence of HGT-markers in different contigs indicates that these MRGs/ARGs might not be highly mobile. However, a few MAGs contained genes annotated as integrative elements (e.g., *intI1*) that can capture genes or are mobile and potentially can be transferred to other locations inside the genome (*tnpA*) and to conjugative plasmids that can transfer to other host taxa (Toussaint and Merlin 2002; Delavat et al. 2017). The occurrence of HGT-markers and ARGs is well described and documented in wastewater or soils from developed countries affected by agricultural activity or pollution (Toussaint and Merlin 2002; Huang et al. 2009; Gatica et al. 2016). Nevertheless, ARG and MRG presence in MAGs assembled from SSA, such as Botswana soils, has been rarely reported. Further studies are needed to understand if these ARGs from bacteria in metal-polluted mines are transferable to bacteria that are common human pathogens or opportunistic pathogens.

### 4.2. *Chloroflexota* adaptation in mine-polluted areas

Collectively, *Chloroflexota* are known for their diverse metabolic capabilities, spanning aerobic and anaerobic conditions, thermophilic adaptability, and anoxygenic photosynthesis (Freches and Fradinho 2024). *Chloroflexota* are known to utilize toxic compounds as electron acceptors, such as As (Wang et al. 2025). This is confirmed by the present study, where we found a significant negative correlation of a single *Chloroflexota* taxon with the distance to the smelter (Fig. 2B). Notably, several taxa belonging to the same phylum positively correlated with distance to the smelter as well. We speculate that *Chloroflexota* might possess an adaptive reservoir and might regulate the transformation of metal or metalloid compounds to a less toxic form. Therefore, our results confirm these previously gained insights that *Chloroflexota* might be helpful for developing a bioremediation strategy for metal-contaminated soils (Wang et al. 2025).

### 4.3. *Actinomycetota* and antimicrobial resistance

*Actinomycetota* (formerly known as *Actinobacteria*) have been considered to possess high bioremediation potential (Alvarez et al. 2017), which was confirmed in the present study since *Actinomycetota* MAGs show several genes annotated as MRGs (Fig. 6). Probably, these bacterial taxa have a high capability for tolerating the high metal concentrations in the mine area, which supports their bioremediation potential. In addition, several *Actinomycetota* taxa can be isolated, and therefore their potential for bioremediation can be confirmed via cultivation-dependent methods. Furthermore, *Actinomycetota* strains are known to produce a diverse amount of antibiotics (Sharrar et al. 2020). They are known to inhabit soils and have been considered to maintain biotic interactions through active competition (Barka et al. 2016). In fact, several antibiotics have been discovered by investigating *Actinomycetota* strains, such as *Actinomyces* spp., for example, several aminoglycosides are known to originate from *Actinomycetota* taxa (van der Meij et al. 2017). Thus, it is not surprising that these bacteria might contain several ARGs to counter the effects of the antibiotic production (Miao and Davies 2010) (Fig. 5). Our findings confirm the occurrence of *Actinomycetota* MAGs, that contain both a high number of MRGs and ARGs in soil from such extreme environments, such as the BCL mine.

### 4.4. *Dormibacterota* and *Acidobacteriota* showed annotated antibiotic resistance and metal resistance genes despite their oligotrophic lifestyle

We here annotated ARGs and MRGs in both *Acidobacteriota* and *Dormibacterota* (Figs 5, 6), which have a small genome size and are known for their oligotrophic lifestyle, where they might depend on other microbial taxa for a nutrient source (Kielak et al. 2016; Sikorski et al. 2022). While *Acidobacteriota* strains have been isolated, *Dormibacterota* have no cultured representative yet (Montgomery et al. 2021). However, cultivation-independent analyses have shown that both *Acidobacteriota* and *Dormibacterota* play an important role in soil biogeochemical functions (Dragone et al. 2024; Gonçalves et al. 2024). So far, little is known about whether these taxa could play a role in bioremediation efforts or AMR dissemination. Therefore, these taxa might also possess a bioremediation mechanism in addition to the presence of MRGs. Moreover, *Acidobacteriota* and *Dormibacterota* MAGs contained genes annotated as HGT-markers despite their low genome size, indicating their potential role in HGT in contaminated soils.

### 4.5. *Thermoproteota* and metal resistance

*Thermoproteota* is a phylum that belongs to *Archaea*, which was previously known as *Crenarchaeota* (Ren and Wang 2022). These archaeal taxa are known for their resistance to heavy metals and salinity tolerance (Li et al. 2023). In the present study, we confirmed their resilience in copper and arsenic contaminated areas. We specifically managed to assemble several *Thermoproteota* MAGs, indicating their high abundance in these soils. In addition, most of these MAGs carried genes annotated as MRGs (Fig. 6), which verify their persistence in the polluted areas. Overall, these results indicate that *Thermoproteota* metabolism could potentially play an important role in the decontamination of toxic metals from the polluted mine.

### 4.6. Limitations of the study

We here analyze MAGs assembled from seven soil metagenomes across a distance to a smelter area. While our results indicate potential selection for ARGs due to metal pollution and co-selection, the limited number of samples did not allow us to conduct a risk analysis. In addition, most of the ARGs were intrinsic ARGs, and not ARGs with high clinical relevance (e.g., resistance to cephalosporins or carbapenems such as *bla*_CTX-M_) or ARGs known to disseminate due to anthropogenic activities (e.g., ARGs that confer resistance to sulfonamides, e.g. *sul1*) (Zhang et al. 2021). This was clear from the fact that annotated ARGs were efflux pumps, which were also annotated as MRGs, due to their low substrate specificity (Pal et al. 2014; Alcock et al. 2023). We consider that further *in situ* experiments in these soils will verify whether metal contamination can co-select for antibiotic resistance or increase the rates of HGT events.

## 5. Conclusions

In this study, we provide for the first time insights into the genomic content of microbes inhabiting Botswana soils near a BCL mine site, an area that is under-sampled, as most areas located in SSA. We show that co-selection of AMR could potentially occur due to metal pollution, using an assembly-based approach and genome-resolved metagenomics. While the co-localization of ARGs and MRGs was very low, many ARGs were also annotated as MRGs, indicating their capability to confer resistance to various substrates. In addition, we describe the potential role of *Actinobacteriota, Chloroflexota, Acidobacteriota,* and *Dormibacterota* in the dissemination of AMR through their resilience in polluted soils. We believe that further sampling and experiments are needed to improve the representation of microbial genomes from under-sampled locations and understand how metal pollution could also lead to the dissemination of AMR in these polluted areas.

## 6. Acknowledgements

This work was supported by de.NBI Cloud within the German Network for Bioinformatics Infrastructure (de.NBI) and ELIXIR-DE (Forschungszentrum Jülich and W-de.NBI-001, W-de.NBI-004, W-de.NBI-008, W-de.NBI-010, W-de.NBI-013, W-de.NBI-014, W-de.NBI-016, W-de.NBI-022).

## 8. Author Contributions

RG and IDK conceived the study. KPPM and OD collected the soil samples. KPPM performed the DNA extraction and analyzed the environmental data. VU assisted with soil analysis and sourcing funding. IDK performed the bioinformatic analysis. DB supervised the analysis. IDK wrote the first draft with inputs from RG and KPPM. All authors contributed to revising the manuscript and approved the final version.

