## Supplementary Information (Legends) for "Genome-resolved metagenomics reveals the potential for selection for antibiotic resistance due to metal pollution in soil microbial communities near a copper-nickel mine site in Botswana"

**Table S1.** Contigs annotated for metal resistance genes (MRGs) using the antibacterial biocide and metal resistance gene database (BacMet). ID= percentage of alignment identity. Query start and end show where the annotation starts and ends for each contig. The sample site of origin is described for each contig as well.

Please look at the corresponding Excel file.

**Table S2.** Contigs annotated for antibiotic resistance genes (ARGs). The annotation was performed based on the comprehensive antibiotic resistance database (CARD). ID= percentage of alignment identity. Query start and end show where the annotation starts and ends for each contig. The sample site of origin is described for each contig as well.

Please look at the corresponding Excel file.

**Table S3.** Contigs annotated for horizontal gene transfer (HGT) markers using the HGT-marker database developed by (https://github.com/KatariinaParnanen/MobileGeneticElementDatabase). ID= percentage of alignment identity. Query start and end show where the annotation starts and ends for each contig. Contigs contain information for their sample of origin as well.

Please look at the corresponding Excel file.

**Table S4.** Metagenome assembled genomes (MAGs) quality characteristics after binning.

Please look at the corresponding Excel file

**Table S5.** Annotated genes as metal resistance genes (MRGs) using the antibacterial biocide and metal resistance gene database (BacMet) from metagenome-assembled genomes (MAGs) after binning.

Please look at the corresponding Excel file

**Table S6.** Annotated genes as antibiotic resistance genes using the comprehensive antibiotic resistance database (CARD) from metagenome-assembled genomes (MAGs) after binning. AMR: antimicrobial resistance.

Please look at the corresponding Excel file

**Table S7.** Annotated genes from the horizontal gene transfer marker database from metagenome-assembled genomes (MAGs) after binning.

Please look at the corresponding Excel file

**Table S8.** MAGs that were annotated for metal-resistance gene (MRGs), antibiotic resistance (ARGs), and horizontal gene transfer markers (HGT-Markers) from metagenome-assembled genomes (MAGs) after binning.

Please look at the corresponding Excel file
